# Basal forebrain parvalbumin neurons modulate vigilant attention

**DOI:** 10.1101/2021.04.19.440515

**Authors:** Felipe L. Schiffino, James M. McNally, Ritchie E. Brown, Robert E. Strecker

## Abstract

Attention is impaired in many neuropsychiatric disorders^1^ and by sleep disruption, leading to decreased workplace productivity and increased risk of accidents^2–4^. Thus, understanding the underlying neural substrates is important for developing treatments. The basal forebrain (BF) is a brain region which degenerates in dementia^5–7^ and is implicated in the negative effects of sleep disruption on vigilance and cognition^8,9^. Previous studies demonstrated that the BF controls cortical fast oscillations that underlie attention^10–12^ and revealed the important role of cholinergic neurons^13–15^. However, the role of other neurochemically defined BF subtypes is unknown. Recent work has shown that one population of BF GABAergic neurons containing the calcium-binding protein parvalbumin (PV) control cortical fast oscillations and arousals from sleep^16–19^ but their role in awake behavior is unclear. Thus, here we test the hypothesis that BF-PV neurons modulate vigilant attention in mice. A lever release version of the rodent psychomotor vigilance test (rPVT) was used to assess vigilant attention as measured by reaction time. Brief and continuous low power optogenetic excitation of BF-PV neurons (1s,473nm@5mW) that preceded the cue light signal by 0.5s improved vigilant attention as indicated by quicker reaction times. In contrast, both sleep deprivation (8h) and optogenetic inhibition of BF-PV neurons (1s,530nm@10mW) slowed reaction times. Importantly, BF-PV excitation rescued the reaction time deficits in sleep deprived mice. These findings reveal for the first time a role for BF-PV neurons in attention.

**HIGHLIGHTS:**

- Optogenetic methods tested the neural circuitry of vigilant attention in mice
- Excitation of basal forebrain parvalbumin neurons quickened reaction times
- Sleep deprivation or inhibition of parvalbumin neurons slowed reaction times
- Excitation of parvalbumin neurons rescued deficits produced by sleep deprivation

## RESULTS

### Assessing vigilant attention in mice using the rodent psychomotor vigilance task (rPVT)

Vigilant attention was assessed in food-restricted mice using a lever release version of the rPVT (**Figure 1A**) which was adapted from previous studies in rats^20–22^. The rPVT is a simple signal detection reaction time test that requires monitoring of a stimulus location (lightbulb) for a brief and unpredictable cue signal (light flash). The lever release rPVT is a self-paced version that assures task engagement by requiring mice to depress a lever to initiate a trial and maintain the lever press until presentation of the cue signal. Importantly, since reaction time is registered as soon as mice release the lever, the measure is not contaminated by movement speed or by locomotion to engage the operandum as in other signal detection task designs^23^. As illustrated in **Figure 1A**, mice must hold the lever down for a random delay (0.7 – 6s) and then report detection of the cue light signal by releasing the lever within the response window (1s) to receive a sucrose pellet reward. Premature responses are defined as lever releases before cue light onset and are punished by lever retraction (3s time-out, no reward delivery). Trials where mice fail to report signal detection within the response window are scored as omissions (i.e., ‘attentional lapses’), no rewards are delivered, and levers remain extended (no time-out). In this study, each rPVT session terminated after 30 rewards were received (session duration typically 10-20 mins). Mice were awake throughout all sessions including those preceded by sleep deprivation.

**Figure 1.**
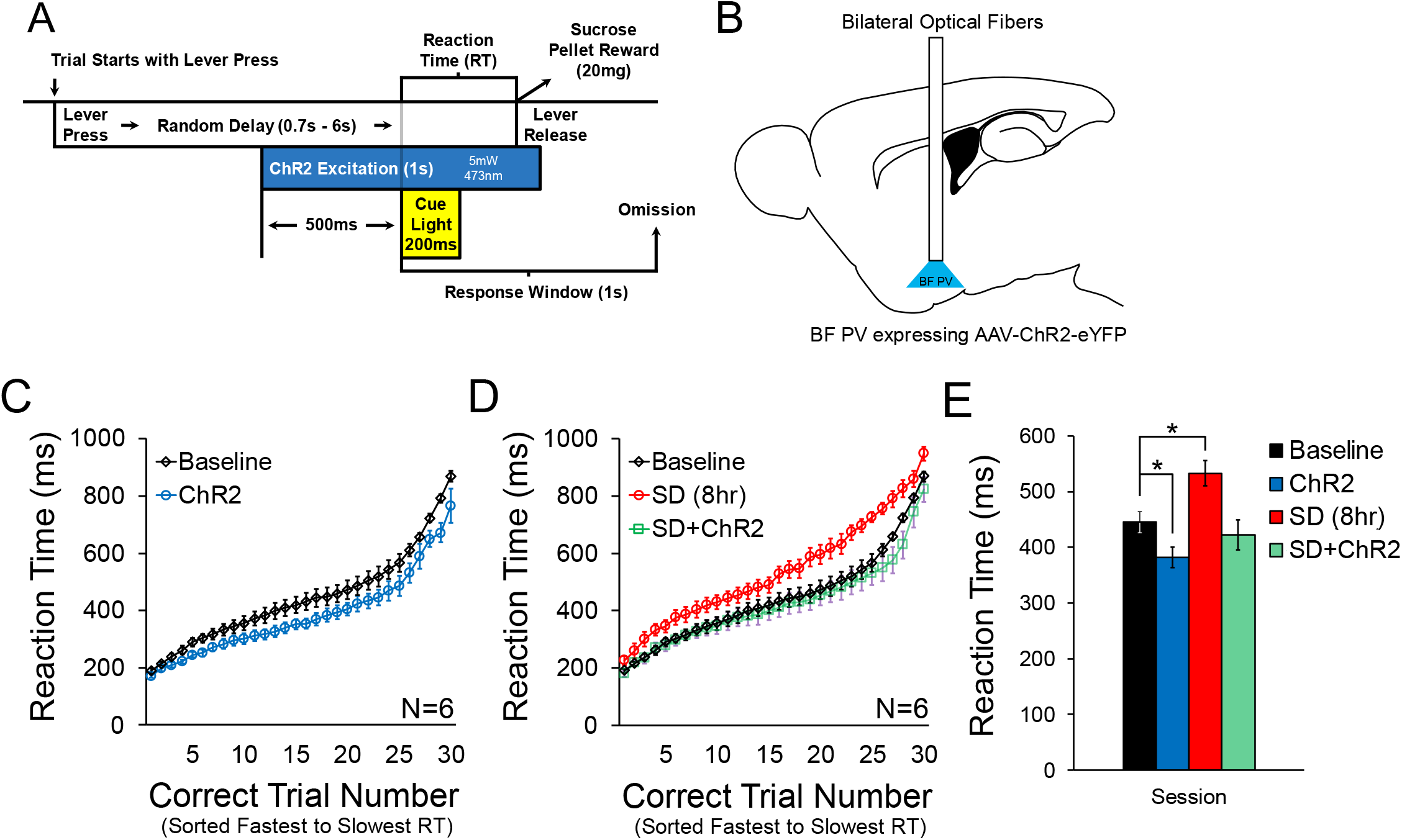
Optogenetic excitation of basal forebrain (BF) parvalbumin (PV) neurons enhances vigilant attention and rescues deficits induced by sleep deprivation (SD). **A:** Design of the lever release version of the rodent psychomotor vigilance task (rPVT) used to assess vigilant attention. Food-restricted mice are required to press a lever down to start each trial. Once the lever is depressed a random delay (0.7– 6.0s) is followed by delivery of a 200ms cue light signal. Mice report cue signal detection by releasing the lever within 1s of signal onset (i.e., a 1s response window) to receive a 20mg sucrose pellet reward for their correct response. Reaction time, the primary measure of vigilant attention, is the time between cue light onset and lever release. Continuous bilateral optogenetic excitation of BF-PV neurons began 500 ms before cue light onset and continued for 500 ms thereafter. **B:** Schematic showing the BF target area for optogenetic excitation. **C:** Optogenetic excitation of BF-PV neurons quickens reaction times (N=6; within-subjects). Sorting correct trial reaction times from fastest to slowest illustrates the consistent pattern of quickened reaction times during BF-PV excitation sessions. **D:** 8 hr of sleep deprivation slows reaction times and optogenetic excitation of BF-PV neurons in sleep deprived mice returns reaction times to baseline levels, indicating rescued performance. **E:** Summary of mean session reaction times in the four experimental groups. Data are represented as mean ± SEM. Asterisk denotes p < 0.05.

### Excitation of BF-PV Neurons Enhances Vigilant Attention and Rescues Deficits Induced by Sleep Deprivation

To test the effect of exciting BF-PV neurons on vigilant attention, we bilaterally injected double-floxed viral vectors into the BF of PV-Cre mice to transduce PV neurons with the excitatory opsin channelrhodopsin2 (ChR2) coupled to enhanced yellow fluorescent protein (EYFP) (**Figure 1B;** STAR Methods). Mice were instrumented with bilateral optical fibers over intermediate/caudal BF (AP +0.4mm, ML ± 1.6 mm, DV −5.2mm) where cortically projecting BF-PV neurons are concentrated^24^. A local field potential electrode was implanted in medial prefrontal cortex to verify efficacy of ChR2 transfection in BF-PV neurons *in vivo* using 40Hz optogenetic BF-PV excitation to evoke 40Hz gamma oscillations as previously shown^16^. Only mice with robust 40 Hz responses were used for experiments. Postmortem histological analysis confirmed that viral expression and fiber placements were localized to the BF (**Figure S1**).

After establishing stable baseline performance (<10% change in mean session reaction time across three consecutive sessions), mice (N=6) were tested under each of three conditions: ChR2 (1s, 5mW continuous BF-PV excitation beginning 0.5s prior to cue light onset; **Figure 1A**), Sleep Deprivation (8hr), and Rescue (sleep deprivation + BF-PV excitation). Bilateral, brief and continuous low-wattage BF-PV excitation was used to enhance activity of BF-PV neurons without attempting to drive specific firing rate frequencies. Moreover, brief excitation closely timed to stimulus onset was used since prolonged (>30s) continuous BF-PV excitation disrupts performance in other tasks^18^.

Sorting trials by reaction time reveals a consistent quickening of reaction time during the BF-PV excitation sessions (**Figures 1C, 1E**). By contrast, sleep deprivation led to a consistent slowing of reaction time, a deficit that was rescued when sleep deprived mice received BF-PV excitation 0.5s prior to the signal light (**Figures 1D, E**). Repeated-measures ANOVA with Greenhouse-Geisser correction on mean session reaction time (**Figure 1E**) shows a significant treatment effect (*F* (3,15) = 37.02, *p* < 0.001) where mean session reaction time is faster in BF-PV excitation sessions relative to baseline (Baseline: 445 ± 19ms vs ChR2: 382 ± 18ms, p < 0.001), slower following 8hr sleep deprivation (SD: 533 ± 23ms, p < 0.01), and returns to baseline when sleep-deprived mice receive BF-PV excitation (SD + ChR2: 422 ± 27ms, p > 0.20) indicating rescued performance. We did not observe significant treatment effects on premature responses (*F* (3,15) = 2.20, *p* > 0.16; Baseline: 33 ± 3; ChR2: 49 ± 8; SD: 29 ± 7; SD + ChR2: 45 ± 8) or on omissions (*F* (3,15) = 3.45, *p* > 0.12; Baseline: 2.3 ± 0.6; ChR2: 1.2 ± 0.4; SD: 9.0 ± 3.8; SD + ChR2: 0.7 ± 0.3). Additional analysis (**Figure S2A**) confirmed that mice responded to the cue light and did not use laser onset to time their response.

### Inhibition of BF-PV Neurons Impairs Vigilant Attention and Mimics the Effects of Sleep Deprivation

To determine the effect of inhibiting BF-PV neurons on vigilant attention, we again used the lever release rPVT task (**Figure 2A**). A viral vector expressing archaerhodopsin (ArchT) / green fluorescent protein (GFP) in the presence of Cre recombinase was bilaterally injected into the BF of PV-Cre mice and bilateral optical fibers were installed in the BF (**Figure 2B**). After establishing stable baseline rPVT performance, mice (N=5) received 1s of continuous 10mW green light for ArchT-mediated BF-PV inhibition beginning 0.5s before cue onset. Sorting trials by reaction time shows that BF-PV inhibition produces a consistent slowing of reaction time (**Figure 2C**). Mean session reaction time (**Figure 2D**) is significantly slower in BF-PV inhibition sessions relative to baseline sessions (Baseline: 426 ± 21ms vs ArchT: 531 ± 11ms, p < 0.02). Although the ArchT mice were not exposed to sleep deprivation, slowing of reaction time by BF-PV inhibition closely resembled reaction time deficits produced by sleep deprivation in the ChR2 mice (c.f., **Figure 1D, E & Figure 2C, D**; SD: 20 ± 4% slower vs. ArchT: 26 ± 7% slower). Finally, BF-PV inhibition increased omissions (Baseline: 2 ± 1 vs ArchT: 14 ± 4, p < 0.05), but did not alter premature responses (Baseline: 39 ± 8 vs ArchT: 31 ± 8, p > 0.58).

**Figure 2.**
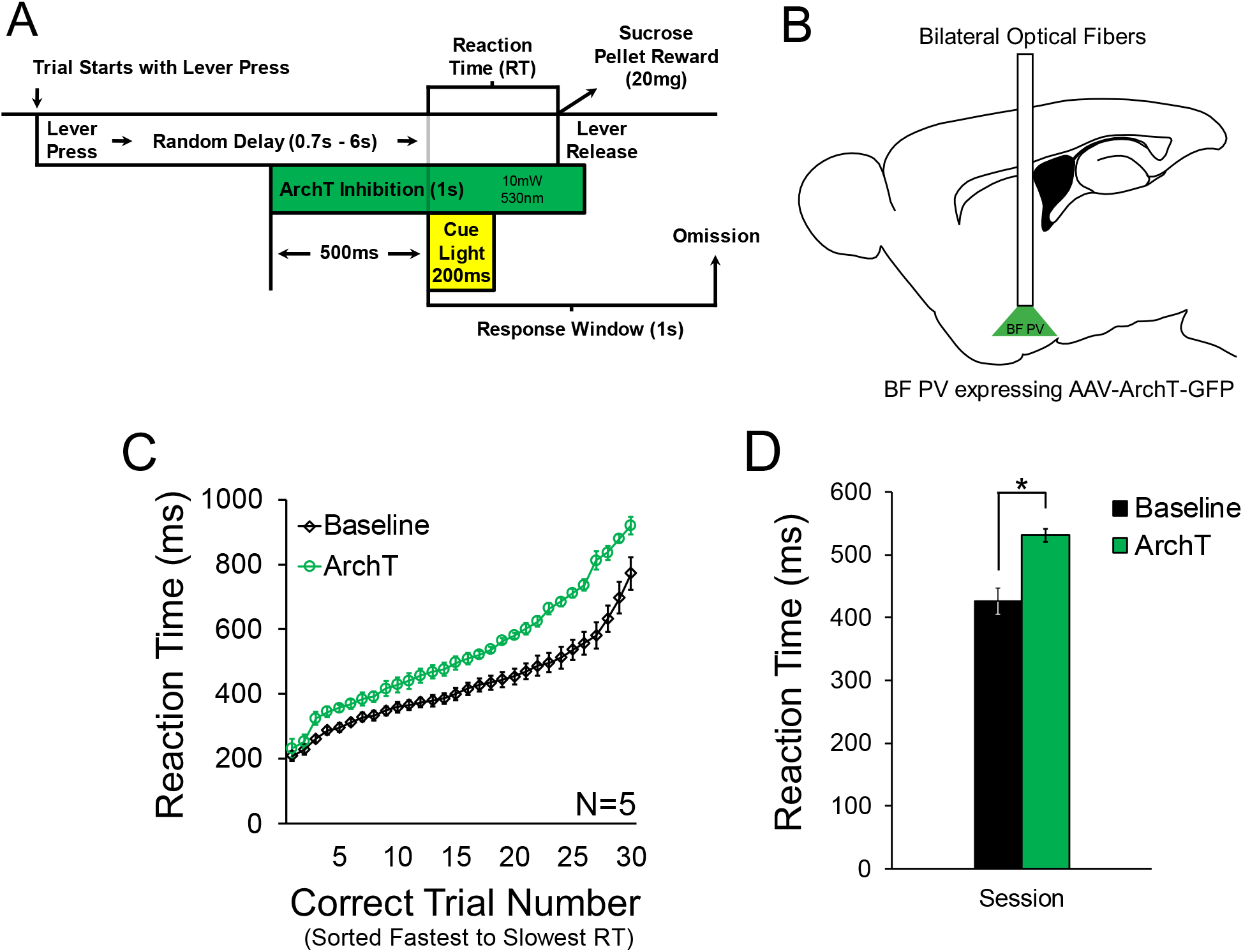
Optogenetic inhibition of BF-PV neurons impairs vigilant attention. **A**. Continuous bilateral optogenetic inhibition of BF-PV neurons began 500 ms before cue light onset and continued for 500 ms thereafter. **B:** Schematic showing the BF target area for optogenetic inhibition. **C:** Sorting correct trial reaction times from fastest to slowest illustrates that BF-PV inhibition (N=5; within-subjects) produces a consistent slowing of reaction times, an effect that closely resembles that of sleep deprivation (see Figure 1). **D:** Mean session reaction time is slower in BF-PV inhibition sessions relative to baseline sessions. Data are represented as mean ± SEM. Asterisk denotes p < 0.05.

### Control Experiments

A group of control mice injected with a non-opsin fluorophore-only encoding control virus (GFP; N=3; **Figure S3**) exhibited no differences in mean session reaction time relative to baseline when given either the blue or green laser light conditions - to control for ChR2 excitation and ArchT inhibition, respectively (*F* (2,4) = 0.19, *p* > 0.98). These control data indicate that altered reaction time during BF-PV excitation/inhibition sessions was not due to non-specific effects of laser illumination. An additional control experiment used the progressive ratio operant task (a method commonly used to assess changes in motivation; e.g. drive to procure food)^25^ to demonstrate that neither BF-PV excitation, nor BF-PV inhibition, altered motivation for food (**Figure S4**).

## DISCUSSION

The BF has long been implicated in the regulation of attention. Neurotoxic lesions of the BF impair attention in rodents^26,27^ and in primates^28,29^. Furthermore, BF degeneration is associated with cognitive deficits, including attention, in Alzheimer’s and Parkinson’s dementias^5–7^. Deficits in attention caused by BF lesions in animals and cognitive deficits associated with dementias have generally been associated with loss of BF cholinergic neurons and their effects in the cortex^13,30–34^. While cholinergic neurons are no doubt important, excitotoxic BF lesions which preferentially affect non-cholinergic neurons have effects on attention which are generally more profound than selective lesions of cholinergic neurons ^26,27,33^. Thus, non-cholinergic BF neurons are likely to be equally important. In fact, single-unit recordings of non-cholinergic neurons, identified by firing rate criteria, have suggested that the activity of several groups of non-cholinergic neurons correlate with attention^35–37^. However, the neurochemical identity of these non-cholinergic, attention-related BF neurons has not been determined.

Several lines of evidence suggested that BF-PV neurons might be one of the BF non-cholinergic subtypes that regulate transient fluctuations in alertness and therefore vigilant attention. BF-PV neurons are large neurons which project to cortex and thalamic reticular nucleus and discharge at high rates (20-60 Hz) during wake^16,24,38^, similar to the discharge rates of a subset of non-cholinergic BF neurons identified in macaques and mice, whose activity ramps up in anticipation of uncertain or unpredictable events^36,37^ and correlates with fast reaction times^36^. Optogenetic excitation of BF-PV neurons rapidly and briefly wakes animals from sleep^19^ and potently enhances fast cortical gamma oscillations^16–18^ which have been linked to successful performance in attention tasks^39^. Furthermore, appropriately timed excitation of BF-PV neurons modulates the cortical topography of sensory stimuli-induced gamma oscillations in a manner reminiscent of the effects of attention^17^. Thus, here we directly tested the role of BF-PV neurons in vigilant attention using optogenetic techniques and a rodent signal detection task that closely mimics human tasks widely used to assess the effects of sleep disruption on attention^40–42^. Optogenetic excitation and inhibition began shortly before delivery of the cue, since unit recording studies showed that a population of fast-firing BF non-cholinergic neurons ramp up their activity after trial start in anticipation of uncertain events^36,37^ in a manner that predicts subsequent reaction time^36^. Moreover, we previously showed that prolonged BF-PV excitation impairs performance in a novel object recognition task^18^. In our experiments here, brief and precisely timed optogenetic excitation of BF-PV neurons led to faster reaction times in the rPVT whereas optogenetic inhibition slowed reaction times, providing the first evidence using gain and loss-of function approaches for a role for BF-PV neurons in attention.

Vigilant attention is altered by the state of arousal and is impaired by sleep loss, fatigue, or other conditions that decrease arousal ^2,40,42^. The BF is one of the most important regions involved in the detrimental effects of sleep loss on cognition^8^. During prolonged wakefulness, the levels of the neuromodulator, adenosine, progressively increase in the BF^43,44^, leading to inhibition of wake-active, cortically-projecting neurons^45,46^. Increased BF adenosinergic tone increases sleepiness^47^ and attentional deficits in a nose-poke version of the rPVT^9^. As with the effects of BF lesions on attention, the role of the BF in sleepiness-induced cognitive deficits has generally been ascribed to inhibition of cholinergic neurons^9,48^. However, *in vitro* patch-clamp experiments showed that adenosine also inhibits the glutamatergic inputs to BF-PV neurons^49^, implicating them in the effects of sleep deprivation. In addition, BF cholinergic neurons innervate and excite BF-PV neurons^50^. Thus, reduced activity of cholinergic neurons likely reduces activity of BF-PV neurons. Here, optogenetic inhibition of BF-PV neurons impaired reaction time in the rPVT to a similar extent as sleep deprivation. Furthermore, we found that optogenetic excitation of BF-PV neurons rescued the effects of sleep deprivation on reaction time, suggesting that direct excitation of these neurons can override the inhibitory effects of adenosine.

A potential confound of our conclusion that BF-PV neurons modulate attention is that excitation or inhibition of BF-PV neurons might affect motivation and therefore alter performance in attention tasks. Indeed, unit recording studies in rodents and macaques have suggested that BF neurons are important for motivationally-guided attention^35,37,51^. However, control experiments here using a progressive ratio operant task suggested that our optogenetic manipulations of BF-PV activity did not affect motivation to procure food. These control experiments are also consistent with findings that optogenetic excitation of BF-PV neurons does not alter food intake^52^. Thus, although the activity of BF-PV neurons may be modulated by inputs from regions regulating motivation^53^, changing their activity does not appear to affect motivation per se.

In a previous study we showed that BF-PV neurons fulfil the criteria for a system which responds to danger signals during sleep by transiently enhancing alertness to support appropriate responses to threat ^19,54^. Our results here show that during wake BF-PV neurons are important for vigilant attention in a food-motivated signal detection task. These new findings support a more general role for BF-PV neurons in upregulating alertness regardless of whether enhanced alertness is motivated by aversive or appetitive situations. Indeed, it is plausible that BF-PV neurons activate cortex in anticipation of, or in response to, motivationally significant events to mobilize cognition and behavior. Thus, therapeutic interventions which enhance the activity of BF-PV neurons may be useful in improving vigilance and therefore attention and cognition in sleep disorders and neurodegenerative diseases.

## AUTHORS’ CONTRIBUTIONS

JMM, REB & RES conceived the experiments. FLS and RES designed the experiments. FLS performed the experiments and analyzed the data. FLS, JMM, REB and RES drafted and revised the manuscript for content.

## ACKNOWLEDGEMENTS

This work was supported by VA Biomedical Laboratory Research and Development Service Merit Awards I01 BX000270 & I01 BX002774 (RES), I01 BX004500 (JMM), I01 BX001356 & I01 BX004673 (REB), VA CDA IK2 BX002130 (JMM) and VA RCS Award IK6 BX005714 (RES); and NIH support from P01 HL095491 (RES), R21 NS079866 (REB), R21 NS093000 (REB), R01 MH039683 (REB), T32 HL07901 (FLS), F32 MH119838 (FLS), and K99 AG066819 (FLS). We thank Drs. Basheer and McKenna for earlier collaborative work that led to this study and helpful discussions, and Abigail Hassler and Leana Radzik for their technical assistance. All authors are scientists at VA Boston Healthcare System, West Roxbury, MA. The contents of this work do not represent the views of the U.S. Department of Veterans Affairs or the United States Government.

## STAR METHODS

### EXPERIMENTAL MODEL AND SUBJECT DETAILS

#### Animals

Adult (4-6 months) homozygous PV-Cre mice (B6.129P2-*Pvalb*^*tm1(cre)Arbr*^/J) were purchased from Jackson Laboratory (Stock#017320; Bar Harbor, Maine) and bred in house. Mice were housed at 21°C with a 12-h light/dark cycle (7AM–7PM light phase), and food and water *ad libitum* until entering behavioral experiment protocols. All procedures were performed in accordance with the National Institutes of Health guidelines and in compliance with the animal protocol approved by the VA Boston Healthcare System Institutional Animal Care and Use Committee.

### METHOD DETAILS

#### Stereotaxic viral vector injections and implantation of optical fibers and LFP/EMG electrodes

For optical excitation of BF-PV neurons we used adeno-associated viral vectors serotype 5 (AAV5), with Cre-recombinase dependent expression of a fusion protein, consisting of ChR2 and EYFP (AAV-DIO-ChR2-EYFP; 3 x 10^12^ viral particles/ml estimated by DotBlot, University of North Carolina Vector Core; Chapel Hill, NC). For inhibition of BF-PV neurons AAV5-CAG-flex-reverse-ArchT-GFP (hereafter referred to as AAV-FLEX-ArchT), bearing the ArchT strain Halorubrum sp. TP009, was purchased from the University of North Carolina vector core. Non-opsin fluorophore-only control experiments used cre-dependent AAV1/2-EGFP vectors (Genedetect, Auckland, NZ) with virion concentration estimated to be 2 x 10^12^ particles/ml by DotBlot. We recently validated BF-PV transduction selectivity and efficiency of these vectors^19^.

Viral vectors (500 nL per side) were bilaterally injected into the BF (AP +0.4mm, ML ± 1.6 mm, DV −5.4mm) of PV-Cre mice under isoflurane anesthesia (induction, 5%; maintenance, 1%–2%) using a Hamilton syringe (50 nl/minute) driven by a high-precision injector pump (model 250; KD Scientific). Optical fibers (200µm, 0.37 NA, Doric Lenses; Quebec City, Quebec, CA) were implanted above BF injection sites (AP +0.4mm, ML ± 1.6 mm, DV −5.2mm) and each mouse was instrumented with an insulated stainless-steel LFP electrode in medial prefrontal cortex (mPFC; AP +1.8mm, ML +0.45mm, DV −1.9-2.0mm). Reference and ground EEG screw electrodes were placed into the skull above the cerebellum on opposite sides of the midline. Electromyography (EMG) electrodes were placed in the nuchal muscle. Electrode leads were connected to EEG/EMG headmounts (Pinnacle Technology Inc., part # 8402-SS, KS, USA) and the installation was secured to the skull using dental cement. The scalp incision was sutured closed and mice were allowed to recover. Effective optogenetic excitation and inhibition of BF-PV neurons using these vectors was previously confirmed by us using *in vitro* electrophysiological methods^16^.

#### Optogenetic Excitation and Inhibition

Bilateral optical illumination was delivered using fiber-coupled 473-nm solid state laser diode (Cat # CL473-050-O; CrystaLaser; Reno, Nevada) or 530-nm laser diode (Cat # CL530-050-O; CrystaLaser; Reno, Nevada) and a patch cord (Doric Lenses) attached to implanted optical fibers with zirconium sleeves. Each sleeve was fitted with black shrink-wrap and the headmount was wrapped with black tape to shield light escape. Software-generated transistor–transistor logic (TTL) pulses (WinWCP; Strathclyde Institute of Pharmacy and Biomedical Sciences, or Spike2; Cambridge Electronic Design) were used to drive optical illumination. After a minimum of four weeks following stereotaxic surgery, mice injected with ChR2-encoding virus were subjected to a 40Hz optogenetic BF-PV excitation protocol to verify efficacy of the manipulation *in vivo*^16^ with 10 ms pulses delivered at 40 Hz (1sON 2sOFF) for 5min (100 trials). LFP signals from medial prefrontal cortex were amplified and conditioned (16 Channel Amplifier; AM System) and recorded with WinWCP at a sampling rate of 2kHz (bandpass filtered at 1-200Hz). Custom MATLAB scripts assessed evoked 40Hz (±5Hz) gamma power. ChR2 animals used for behavioral experiments exhibited >2000 µV^2^/Hz evoked 40Hz (±5Hz) gamma power during photoexcitation. For BF-PV optogenetic manipulations during rPVT performance, 1s of continuous laser illumination (5mW 473nm blue light for ChR2-mediated BF-PV excitation; 10mW 530nm green light for ArchT-mediated BF-PV inhibition) was delivered on every trial beginning 500ms prior to cue light onset. Animals injected with non-opsin encoding fluorophore control virus (GFP) received delivery of blue and green laser in separate sessions.

#### Behavioral procedures

Mice were subjected to one behavioral session per day during the light phase (7am-7pm). During the post-surgical recovery period (at least one week), mice were individually housed and provided with *ad libitum* access to water and rodent chow. Mice were then food-restricted and fed 2-3g of chow per day to maintain 90% of their *ad libitum* weight throughout experiments. To acclimate to the food rewards (20 mg sucrose pellets, TestDiets 5TUT), mice were given 10 pellets/day in their home cages for three days prior to the onset of behavioral training. Both behavioral procedures (rodent psychomotor vigilance task; rPVT; progressive ratio task, PR) consisted of transferring mice to software-controlled operant chambers (GraphicState4.0; Coulbourn Instruments LLC, PA, USA) equipped with two retractable levers on either side of a sucrose pellet dispenser and panel lights located directly above each lever. In both experiments, mice were first trained to retrieve rewards from the dispenser through a pseudorandom delivery of pellets (30 pellets over a 60 min session). In the next session, mice were trained to press the active lever to receive a single pellet per press (fixed ratio; FR1) until 30 pellets were earned. Presses of the inactive lever had no programmed consequence and lever designations were counterbalanced across mice. The procedures for the two experiments then diverged.

In the rPVT experiment, the next six sessions trained mice to press and hold the active lever down for an increasing amount of time (hold duration) to receive pellet rewards (30 pellets per session). The hold duration criterion began at 200ms for each of the 30 rewards in the first session. In the next session, hold criterion remained at 200ms for the first five rewards, increased to 260ms for the next ten rewards, then increased to 300ms for the last 15 rewards. Incremental training continued in this manner across the next four sessions until a final hold duration of 700ms was achieved. The next phase of the rPVT experiment trained mice to release the lever in response to brief illumination of the panel light cue (200ms) located directly above the active lever. First, mice were introduced to the pseudorandom presentation of the light signal after a minimum hold of 500ms. Then, mice were required to release the lever within 1s of cue light onset (response window) to receive reward (correct trial) while premature responses (lever release before cue onset) were punished by retraction of both levers for 3s (time-out). If mice failed to release the lever within 1s after light onset, the trial was scored as an omission (lapse in attention), no reward was delivered, and the levers remained extended (no time-out). The time between cue onset and lever release on correct trials is the primary measure of vigilant attention (reaction time: RT). Training continued until performance was stable across 3 sessions (<10% Δ in mean session RT) to establish a baseline for each mouse. Sleep deprivation was achieved by gentle handling for 8hr prior to rPVT sessions.

In the progressive ratio experiment, the FR1 session was followed by two sessions of FR3 (three presses per pellet). Then, mice were trained on a progressive ratio schedule according to the following formula: [Response Requirement = (5*e*^0.2 × Pellet Number^) – 5] as in refs^25,57^, until the number of pellets earned was stable across three sessions (<10% change). Optogenetic manipulations of BF-PV activity used the same parameters as above and delivered laser for 1s every 30s throughout the 1hr test session. The measure of motivation is the number of pellets earned.

#### Perfusion/brain extraction

Mice were anesthetized with sodium pentobarbital (50 mg/ml), exsanguinated with ice-cold PBS, and perfused transcardially with 10% formalin (Cat # HT5011; Sigma-Aldrich; St. Louis, MO). Brains were post-fixed overnight in 10% formalin then transferred to 30% sucrose solution. 40 mm-thick coronal slices were collected and stored at 4°C.

#### Tissue mounting/microscopy and photography

Coronal sections were mounted onto gel-alum subbed slides and coverslipped using Vectashield Hard Set mounting medium (Cat #H-1400; Vector Laboratories; Burlingame, CA). Fluorescent microscopy and photography were performed using a Zeiss Image2 microscope, with a Hamamatsu Orca R2 camera (C10600). Optogenetic fiber optic cannula locations were mapped onto appropriate schematic templates employing Adobe Illustrator (v.CS5.1).

