## Supplemental Figures for "Basal forebrain parvalbumin neurons modulate vigilant attention"

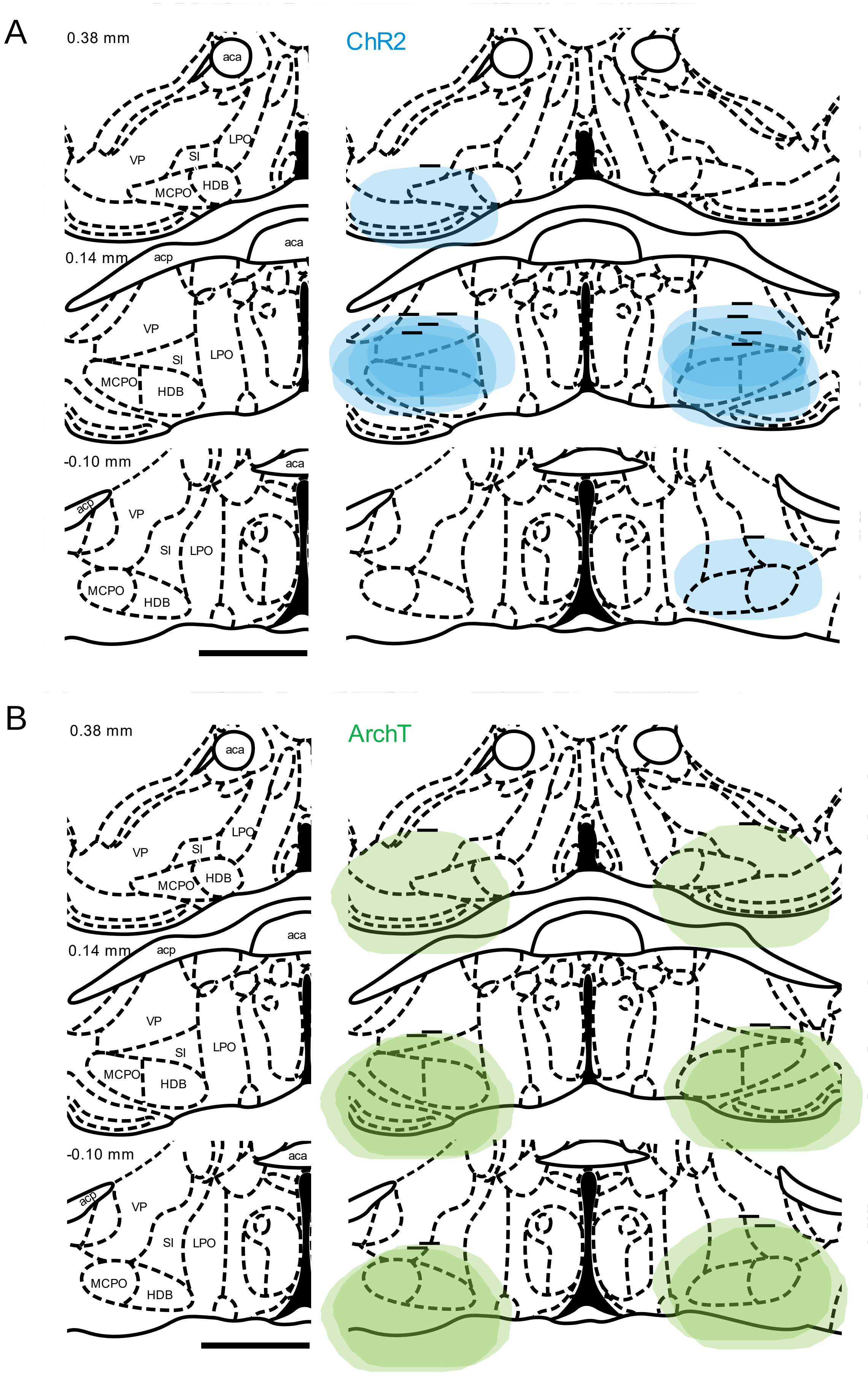
A
0.38 mm
aca
LPO
VP
SI
MCPO
HDB
ChR2
0.14 mm
acp
aca
VP
SI
LPO
MCPO
HDB
-0.10 mm
aca
acp
VP
SI
LPO
MCPO
HDB
B
0.38 mm
aca
LPO
VP
SI
MCPO
HDB
ArchT
0.14 mm
acp
aca
VP
SI
LPO
MCPO
HDB
-0.10 mm
aca
acp
VP
SI
LPO
MCPO
HDB

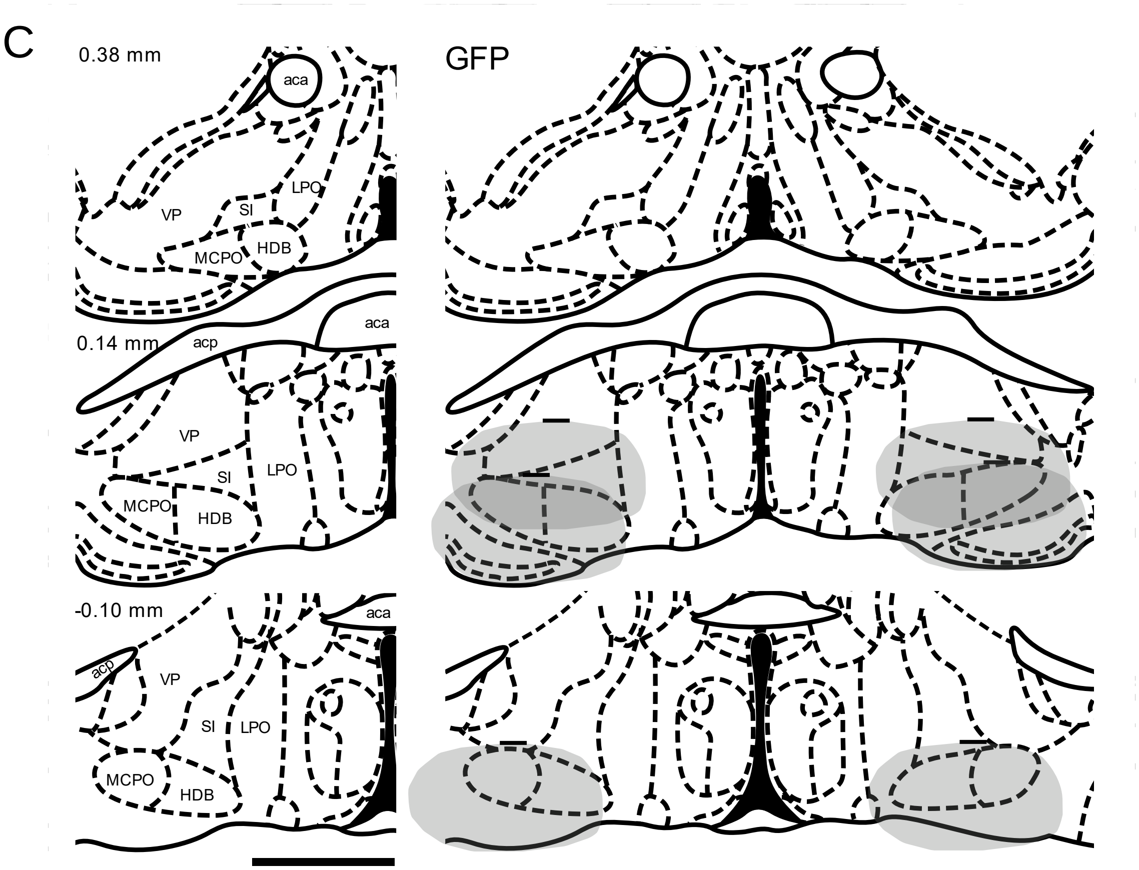


**Supplemental Figure 1. Summary of location of histologically confirmed optical fiber tips and theoretical area of laser irradiance > 1mW/mm$^2$.** Localization of bilateral optogenetic fiber tips (horizontal black lines) in BF for **A:** ChR2-mediated BF-PV excitation ('ChR2'; N=6, blue), **B:** ArchT-mediated BF-PV inhibition ('ArchT'; N=5, green), and **C:** non-opsin fluorophore control ('GFP'; N=3, gray) mice used in this study. Contours for theoretical area of irradiance > 1mW/mm$^2$ were estimated using the Deisseroth laboratory on-line calculator (https://web.stanford.edu/group/dlab/cgi-bin/graph/chart.php) and depictions of scattered light in rodent gray matter in Fig. 3E of Yizhar et al., (2011)[55]. Scale bar = 1mm. Abbreviations: aca, anterior aspect of the anterior commissure; acp, posterior aspect of the anterior commissure; HDB, horizontal limb of the diagonal band; LPO, lateral preoptic nucleus; MCPO, magnocellular preoptic nucleus; SI, substantia innominata; VP, ventral pallidum. Atlas templates adapted from Paxinos & Watson (2008)[56].

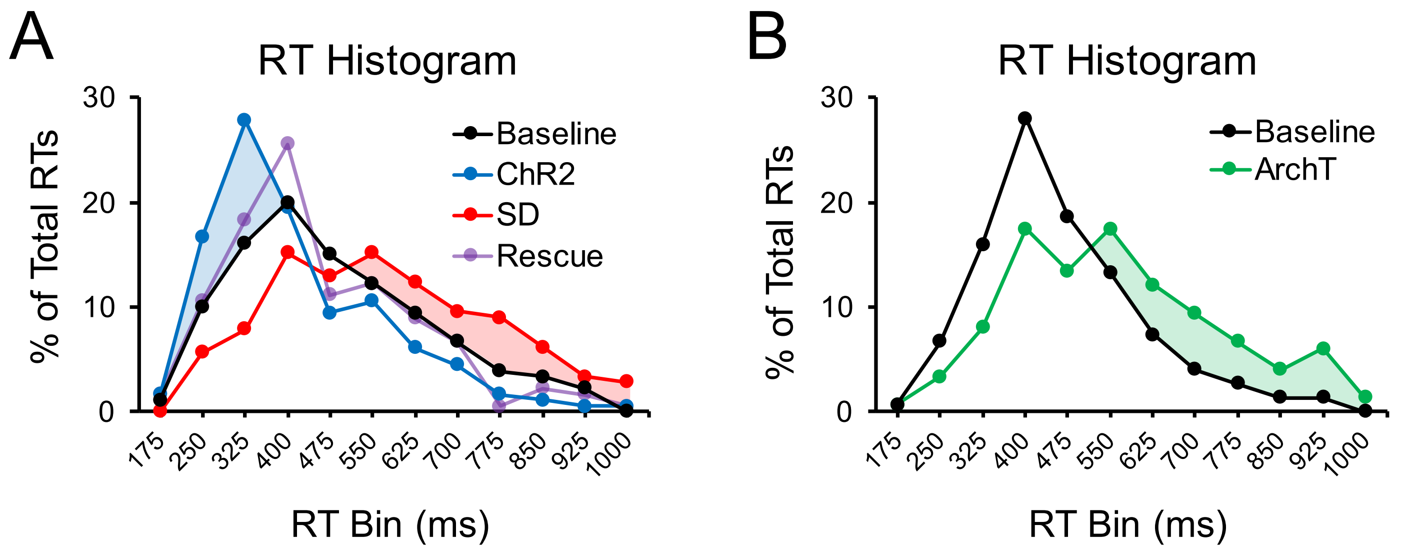


**Supplemental Figure 2. Reaction time (RT) distributions show that BF PV excitation increases the frequency of quicker RTs within the physiological range and that BF PV inhibition slows RT in a manner similar to the effect of sleep deprivation (SD).** This figure plots RT frequency histograms in 75ms bins (12 bins total spanning RTs < 175ms to RTs between 925-1000ms, the upper limit of each bin is demarked on the *x*-axis) to further characterize the effects of our manipulations and show that quickened RTs during BF-PV excitation sessions is not due to mice using the laser onset to time their response. **A:** Analysis of RT histograms for the BF-PV excitation experiment indicates that during baseline performance without laser, a majority of correct trials registered RTs between 175-475ms (61% of total responses after collapsing across the four bins) with a plurality between 325-400ms (20% for that bin), a considerable number of ‘fast’ responses between 175-325ms (26% across the two bins), and very few trials registering RT < 175ms (1%). Importantly, BF-PV excitation did not appreciably increase the frequency of trials with RT < 175ms (2%) as would be expected if mice were timing cue light onset from laser onset. Instead, BF-PV excitation accelerated RT such that most correct trials registered RTs between 175-400ms (64% across three bins) with a plurality between 250-325ms (28%) and an increased number of ‘fast’ RTs within the physiological range of 175-325ms (44% across two bins). SD widens the distribution of RTs reducing the number of faster RTs (<400ms; 36 ± 9% decrease) and increasing the number of slow RTs (>550ms; 84 ± 41% increase). The RT distribution following BF-PV excitation in sleep deprived animals closely approximates the baseline distribution indicating rescued performance. **B:** BF-PV inhibition widens the RT distribution like SD by reducing fast RTs and increasing slow RTs.

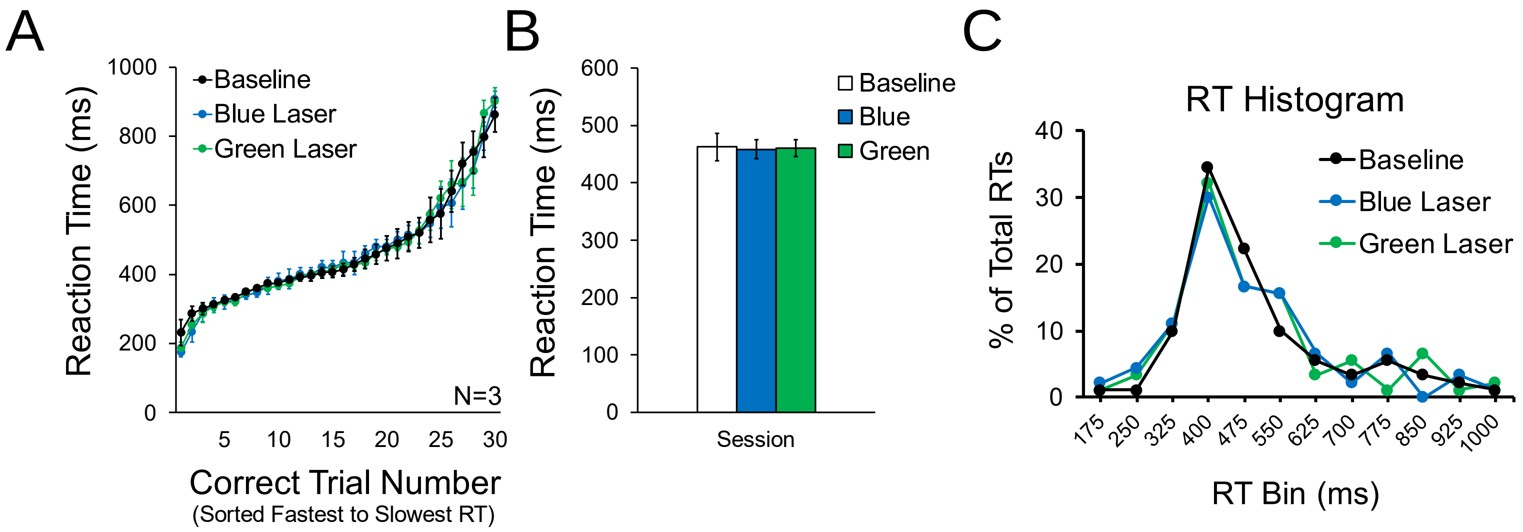


**Supplemental Figure 3. In non-opsin control mice vigilant attention is not altered by blue or green laser delivery.** Mice expressing only non-opsin green fluorescent protein (GFP) control virus in BF-PV neurons (N=3; within-subjects) exhibit no differences in mean session RT in the rPVT relative to baseline when given the blue or green laser delivery parameters used in the ChR2 and ArchT experiments ($F$ (2,4) = 0.19, $p$ > 0.98). These data provide additional evidence that: i) in BF-PV excitation sessions the quicker RTs are not due to mice using laser onset to predict cue light onset and, ii) in BF-PV inhibition sessions that slower RTs are not due to potential distracting effects of green laser.

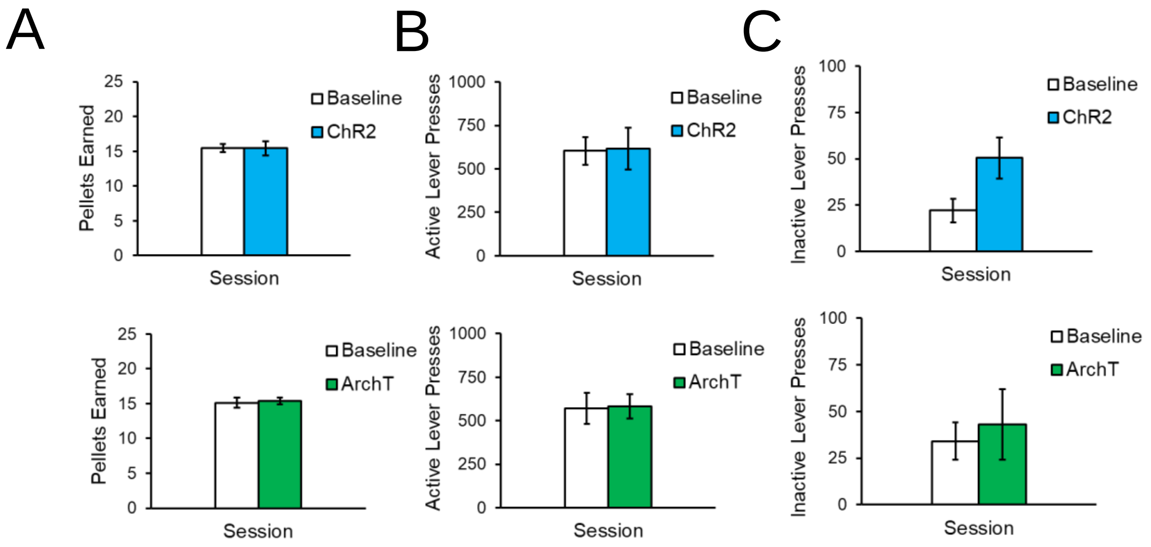


**Supplemental Figure 4. Optogenetic manipulations of BF-PV neuronal activity do not alter motivation for food as assessed by a progressive ratio operant task.** The progressive ratio task is commonly used to assess motivation (e.g. drive to procure food). It does so by increasing the effort required for each successive reward. Highly motivated subjects will expend greater effort to receive more rewards. The primary metric of motivation therefore is the total number of sucrose pellets earned. The response requirement (lever press) for rewards was calculated according to the following formula: [Response Requirement = $(5e^{0.2 \times \text{Pellet Number}}) - 5$] (Richardson & Roberts, 1996; Schiffino et al., 2019)[25,57]. Mice were trained on the task until the number of pellets earned was stable across three consecutive sessions (<10% change). BF-PV excitation (5mW continuous blue laser) or BF-PV inhibition (10mW continuous green laser) was delivered for 1s every 30s throughout the 1hr session to ascertain whether manipulation of BF-PV activity alters motivation for food.

(A) Neither BF-PV excitation (N = 7, top) nor BF-PV inhibition (N = 5, bottom) altered the total number of pellets earned nor

(B) the total number of active lever presses.

(C) BF-PV excitation increased the number of inactive lever presses (Baseline: 22 ± 6 vs ChR2: 50 ± 10; $p < 0.005$) indicating slight hyperactivity, which is consistent with a role for BF-PV neurons in arousal.
